# Harnessing a Novel P450 Fatty Acid Decarboxylase from *Macrococcus caseolyticus* for Microbial Biosynthesis of Odd Chain Terminal Alkenes

**DOI:** 10.1101/255539

**Authors:** Jong-won Lee, Narayan P. Niraula, Cong T. Trinh

**Author notes:** Equal contributions (Jong-won Lee, Narayan P. Niraula). Current address: Pfizer Inc., Kalamazoo, MI.

## Abstract

Alkenes are industrially important platform chemicals with broad applications. In this study, we report a microbial conversion route for direct biosynthesis of medium and long chain terminal alkenes from fermentable sugars by harnessing a novel P450 fatty acid (FA) decarboxylase from *Macrococcus caseolyticus* (OleT_MC_). We first characterized OleT_MC_ and demonstrated its *in vitro* H_2_O_2_-independent activities towards linear and saturated C10:0-C18:0 FAs, with the highest activity for C16:0 and C18:0 FAs. Combining protein homology modeling, *in silico* residue mutation analysis, and docking simulation with direct experimental evidence, we elucidated the underlying mechanism for governing the observed substrate preference of OleT_MC_, which depends on the size of FA binding pocket, not the catalytic site. Next, we engineered the terminal alkene biosynthesis pathway, consisting of an engineered *E. coli* thioesterase (TesA*) and OleT_MC_, and introduced this pathway into *E. coli* for direct terminal alkene biosynthesis from glucose. The recombinant strain *E. coli* EcNN101 produced a total of 17.78 ± 0.63 mg/L odd-chain terminal alkenes, comprising of 0.9% ± 0.5% C11 alkene, 12.7% ± 2.2% C13 alkene, 82.7% ± 1.7% C15 alkene, and 3.7% ± 0.8% C17 alkene, and a yield of 0.87 ± 0.03 (mg/g) on glucose after 48 h in baffled shake flasks. To improve the terminal alkene production, we identified and overcame the electron transfer limitation in OleT_MC_, by introducing a two-component redox system, consisting of a putidaredoxin reductase CamA and a putidaredoxin CamB from *Pseudomonas putida,* into EcNN101, and demonstrated the terminal alkene production increased ∼2.8 fold after 48 h. Overall, this study provides a better understanding of the function of P450 FA decarboxylases and helps guide future protein and metabolic engineering for enhanced microbial production of target designer alkenes in a recombinant host.

## 1. INTRODUCTION

Alkenes (or olefins) are industrially important platform chemicals used to manufacture polymers, lubricants, surfactants, and coatings (1). Alkenes are currently produced by the well-established chemical conversion route (e.g., hydrogen cracking) using petroleum-based feedstocks that are neither renewable nor sustainable (2, 3). In recent years, there is great interest in developing microbial conversion routes to produce alkenes from renewable and sustainable sources, such as biomass-derived fermentable sugars (4–10).

Various species are known to produce alkenes endogenously via decarbonylation of aldehydes (11–13), decarboxylation of FAs (14), condensation of unsaturated polyenes (15), and oxidation of FAs (7, 8). Various types of alkenes can be synthesized including alkadienes (16, 17), terminal alkenes (or 1-alkenes) (4, 6–8) as well as non-terminal alkenes (5), depending on enzyme types and substrates employed. To date, a number of different classes of enzymes have been reported to synthesize terminal alkenes including a P450 FA decarboxylase/peroxygenase (OleT, belonging to the CYP152 family) (4), a type-I polyketide synthase-like enzyme (CurM/Ols) (6), a desaturase-like enzyme (UndB) (7), and a non-heme oxidase (UndA) (8). These enzymes take various substrates (e.g., FAs, fatty aldehydes, or FA thioesters) to produce terminal alkenes with different carbon chain lengths. For instance, OleT, CurM/Ols, and UndB are capable of synthesizing (C_6_-C_12_) medium-chain and (> C_12_) long-chain alkenes while UndA only produces medium chain length alkenes. The diversity of these enzyme specificities can potentially offer unique opportunities to develop microbial cell factories to engineer designer olefins for tailored applications.

Among the putative P450 FA decarboxylases/peroxygenases discovered, OleT_JE_ is the most well-characterized enzyme that is capable of using either H_2_O_2_−dependent or H_2_O_2_−independent (O_2_-dependent) mechanisms for alkene biosynthesis (4, 16, 18, 19). OleT_JE_ has broad substrate specificity for C12:0-C20:0 FAs *in vitro*, with the highest towards C12:0 FA in the presence of redox partner proteins and C14:0 FA in the H_2_O_2_−dependent system (16). For the low-cost, large-scale production of terminal alkenes, the use of H_2_O_2_-independent decarboxylases (e.g., OleT_JE_) is likely favorable by avoiding the external supply and cytotoxicity of H_2_O_2_. Thus, harnessing the enzyme library of H_2_O_2_−independent decarboxylases are important for the microbial production of terminal alkenes from biomass-derived sugars.

In this study, we expanded the library of unique OleTs for terminal alkene biosynthesis by characterizing a novel P450 FA decarboxylase OleT_MC_ derived from *Macrococcus caseolyticus* for its catalytic functions. We employed protein homology modeling, *in silico* residue mutation analysis, and docking simulation to elucidate the underlying mechanism of fatty acid specificity of OleT_MC_. Furthermore, we harnessed this OleT_MC_ to engineer a terminal alkene biosynthesis pathway in *E. coli* for direct conversion of fermentable sugars into medium and long chain terminal alkenes (Figure 1). We discovered that electron transfer was the rate limiting step of alkene biosynthesis in the recombinant *E. coli*. This limitation could be alleviated by expressing the redox system CamA/B from *Pseudomonas putida* in the recombinant *E. coli*, resulting in enhanced alkene production.

**Figure 1:**
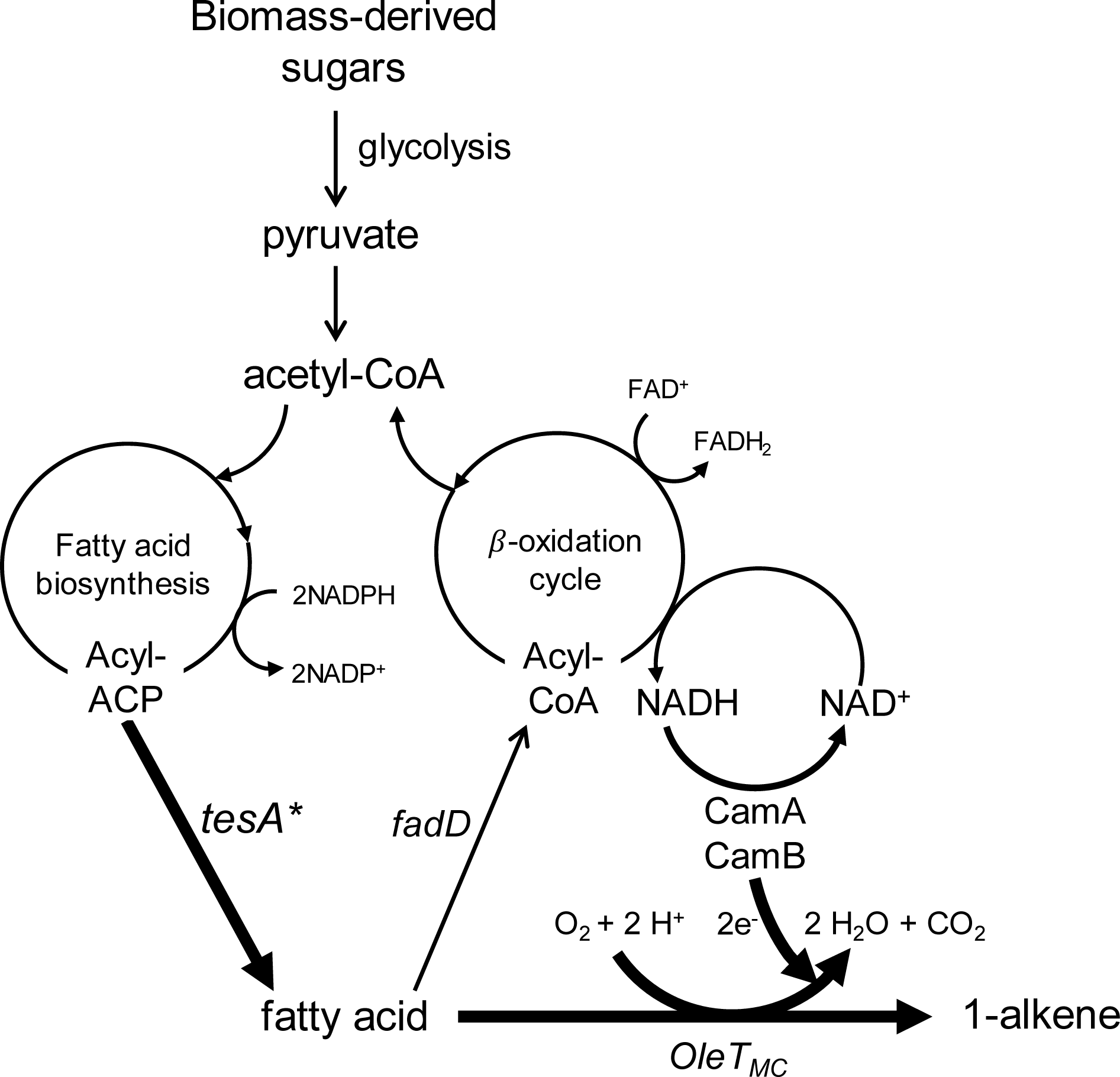
Synthetic pathway for endogenous production of terminal alkenes in *E. coli*

## 2. RESULTS AND DISCUSSION

### 2.1 Genome mining of OleT decarboxylases for terminal alkene biosynthesis

To identify the putative H_2_O_2_-independent P450 FA decarboxylases, we performed a genome mining, a combination of the sequence alignment and phylogenetic analysis, using the protein sequence of OleT_JE_ (ADW41779) as a template. The sequences of the CYP152 P450 enzyme family (4) were first aligned to select the candidates that have the conserved catalytic site residues Phe79, His85, and Arg245 like OleT_JE_ (20). Among the 29 decarboxylase candidates, we found that three P450 enzymes from *M. caseolyticus* (WP_041635889.1), *Corynebacterium efficiens* (WP_011075937.1), and *Kocuria rhizophila* (WP_012399225.1) have the conserved catalytic site residues (Supplementary Figure S1). Based on the phylogenetic analysis, we found that the P450 enzyme of *M. caseolyticus* (WP_041635889.1) is the closest ortholog to OleT_JE_ with the highest amino acid identity (∼60%) (Figure 2 and Supplementary Figure S1). Thus, we chose the P450 from *M. caseolyticus,* named OleT_MC_, for further characterization.

**Figure 2:**
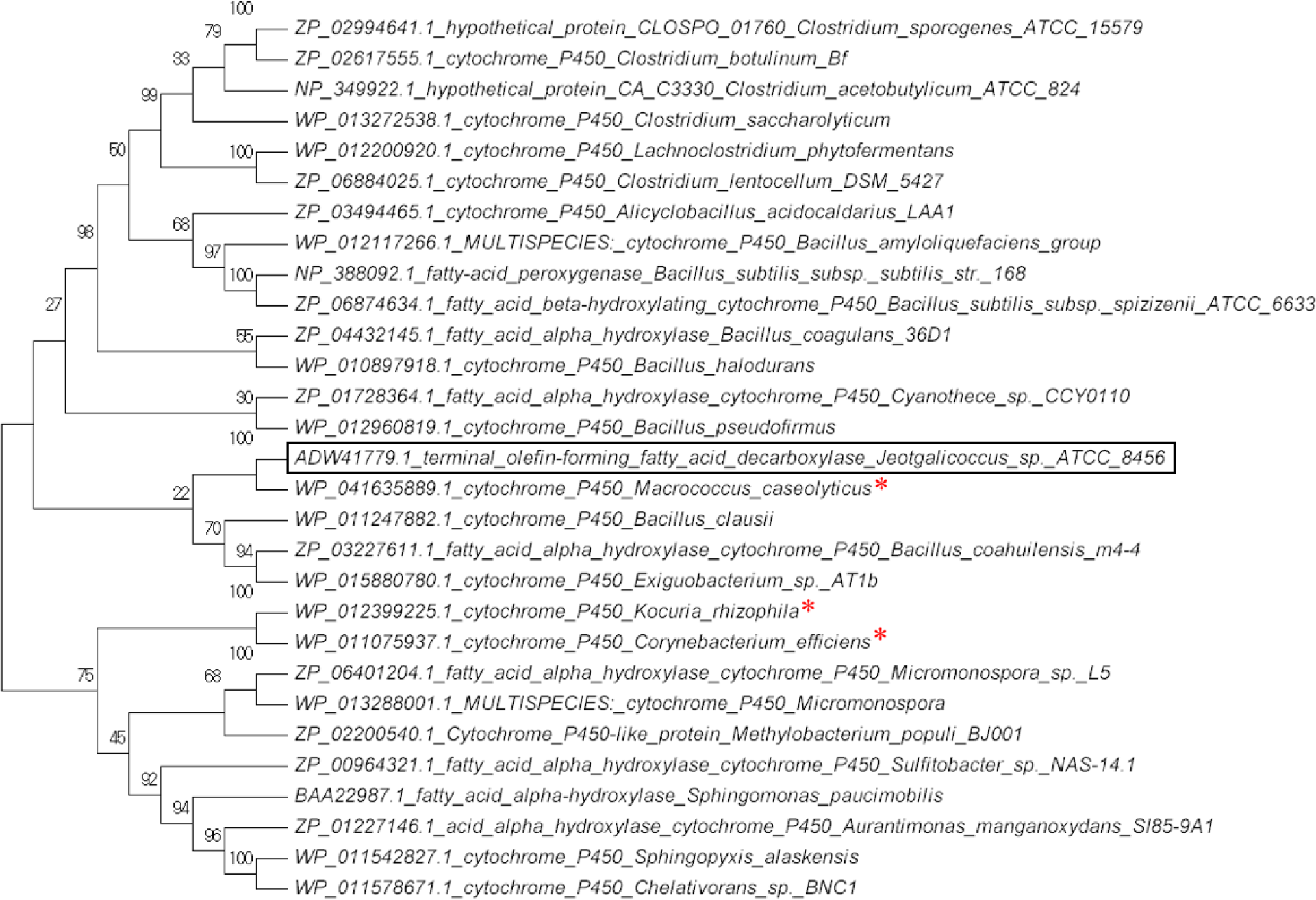
Phylogenetic analysis of OleT_JE_ with the CYP152 P450 enzyme family. OleT_JE_ is shown in the box. The enzymes that have the conserved OleT_JE_ catalytic site residues Phe79, His85, and Arg245 are marked with “*”

### 2.2 *In vitro* characterization of OleT_MC_

The expression of OleT_MC_ in BL21 (DE3) pNN33 was confirmed *in vivo* with reddish cell cultures due to the heme-containing OleT_MC_ and *in vitro* by a sodium dodecyl sulfate polyacrylamide gel electrophoresis (SDS-PAGE) and spectrophotometric analysis (Supplementary Figure S2). After protein isolation, we performed the *in vitro* enzyme assay to examine the H_2_O_2_-independent decarboxylase activity of OleT_MC_ towards linear, saturated FAs. The result shows that OleT_MC_ could convert C10:0-C18:0 FAs to the corresponding odd chain terminal alkenes without H_2_O_2_ as an oxidant, confirmed by GC/MS (Supplementary Figure S3). Under the H_2_O_2_-independent (O_2_-denpendent) conditions, OleT_MC_ showed the highest specific activity towards C18:0 FA (1.13 ± 0.09 μM/min/mg) and the lowest specific activity towards C10:0 FA (0.17 ± 0.04 μM/min/mg) (Figure 3A). OleT_MC_ exhibited almost the same specific activity for C12:0 FA (0.62 ± 0.10 μM/min/mg) and C14:0 FA (0.66 ± 0.04 μM/min/mg). We did not observe the activity of OleT_MC_ for < C10:0 FAs. The FA specificity of OleT_MC_ can be ranked as follows: C18:0 > C16:0 > C14:0 ≥ C12:0 > C10:0. Taken altogether, OleT_MC_ is a potential FA decarboxylase for developing the terminal alkene biosynthesis pathway in recombinant hosts (e.g., *E. coli*) for direct conversion of fermentable sugars to terminal alkenes.

**Figure 3:**
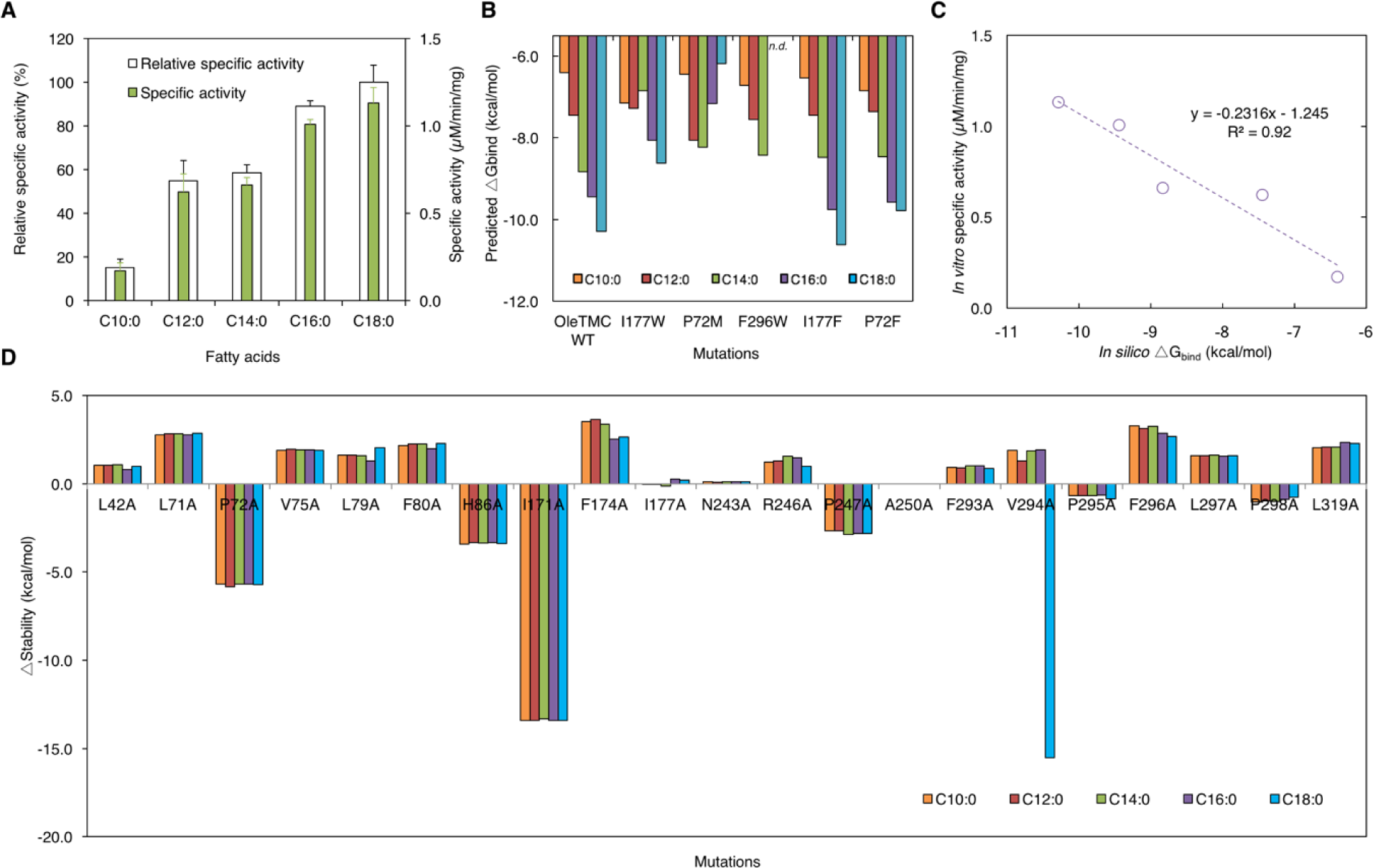
**(A)** Specific and relative specific activities of OleT_MC_ towards linear, saturated C10:0-C18:0 FAs. Each value is an average ± 1 standard deviation (n ≥ 2). **(B)** Comparison of predicted binding free energies for C10:0-C18:0 FAs between the OleT_MC_ wildtype and mutants. **(C)** Correlation between *in vitro* specific activities and *in silico* binding free energies of OleT_MC_ with various C10:0-C18:0 FAs. **(D)** *In silico* alanine scan of the potential FAs binding pocket with C10:0-C18:0 FAs-OleT_MC_ complexes.

### 2.3 Elucidate the underlying mechanism governing the substrate specificity of OleT_MC_

Currently, it is not well understood why different FA decarboxylases have different substrate specificities. For instance, OleT_JE_ prefers C12:0-C14:0 FAs to longer ones (16, 20) while OleT_MC_ favors C16:0-C18:0 FAs more than shorter ones (this study). To better understand the underlying mechanism for governing the observed substrate preference of OleT_MC_, we focused on analyzing its protein structure, using protein homology modeling, *in silico* residue mutation analysis, and docking simulation in combination with direct experimental evidence.

#### 2.3.1 Analysis of the effect of catalytic site on the substrate specificity of OleT_MC_

Since OleT_MC_ and OleT_JE_ have different substrate preferences, we first tested whether structural difference of their catalytic sites might exist and play a role. To do this, we built a 3D structure of OleT_MC_ using the version 2015.10 Molecular Operating Environment (MOE) software (21), based on the best-hit crystallographic structure of the substrate-bound (C20:0, arachidic acid) form of OleT_JE_ (PDB:4L40 (19), ∼60% amino acid identity) as a template. The Ramachandran plot of OleT_MC_ showed less than 0.5% of the residues to be in disallowed regions (Supplementary Figure S4). Next, we superposed the heme-bound 3D structures of OleT_MC_ and OleT_JE_ and found that they look almost identical (Figure 4A) with a very similar catalytic site configuration (Figure 4B). This result suggests that other factors, not the catalytic site, might be responsible for modulating the substrate specificities of OleT_MC_ and OleT_JE_. This observation concurs with recent experimental evidence that single site directed mutations on the catalytic site residues, including Phe79, His85, and Arg245, did not improve the catalytic activity or change the substrate preference of OleT_JE_ (20, 22, 23). Further, a recent molecular dynamics study suggests several residues of OleT_JE_ interacting with FA could significantly contribute to the substrate binding free energy (24).

**Figure 4:**
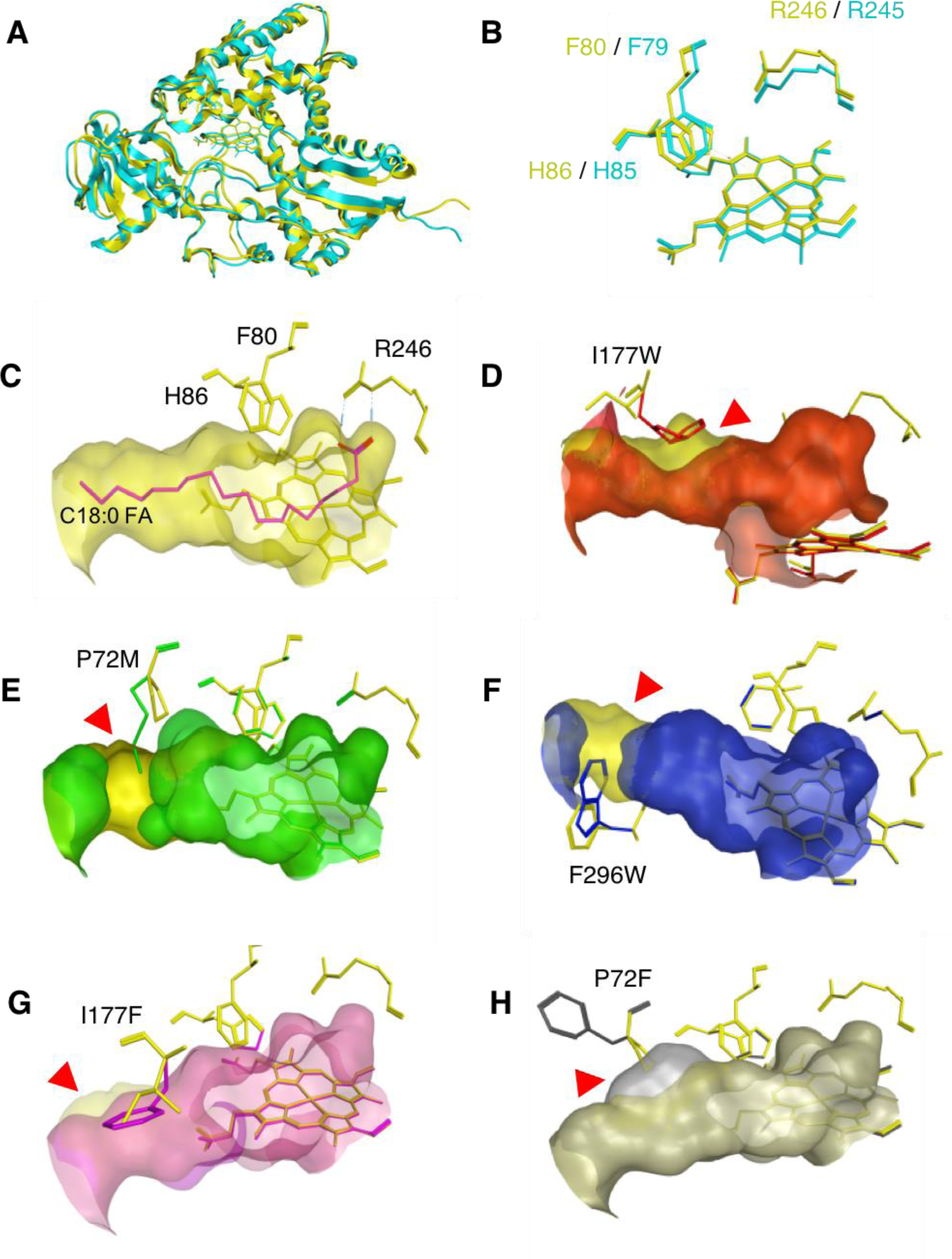
**(A)** Comparison of the protein structures between the homology model of OleT_MC_ (in yellow) and the crystal structure of C20:0 FA-bound OleT_JE_ (in cyan; PDB:4L40). **(B)** Overlay of the catalytic site structures of OleT_MC_ (in yellow) and OleT_JE_ (in cyan). **(C)** A homology model of OleT_MC_ docked with C18:0 FA. **(D-H)** Overlay of the potential FA binding pocket of the wildtype OleT_MC_ (in yellow) and the OleT_MC_ variants including **(D)** I177W (in red) **(E)** P72M (in blue) **(F)** F296W (in purple) **(G)** I177F (in pink) **(H)** P72F (in gray). Filled red triangles point to the distinctive features in the FA binding pockets of the OleT_MC_ variants as compared to the wildtype.

#### 2.3.2 Effect of binding free energy of OleT_MC_ on its substrate preferences

To examine whether the binding free energy of FA-bound OleT_MC_ might affect its substrate specificity, we docked OleT_MC_ with various FAs (C10:0-C18:0) in MOE. Our result shows that the FA-bound OleT_MC_ complexes exhibited the interactions between FAs and OleT_MC_ where the binding pocket of OleT_MC_ contains greasy residues such as alanine (Ala, A), valine (Val, V), leucine (Leu, L), isoleucine (Ile, I), proline (Pro, P), phenylalanine (Phe, F) (Supplementary Figures S5 and S6). The binding free energies of different FA-bound OleT_MC_ complexes increase with shorter chain FAs (Figure 3B). Remarkably, there is a strong linear correlation (R^2^ = 0.92) between the binding free energies and specific activities of *OleT_MC_* for various FAs (Figure 3C).

#### 2.3.3 Residues at the tail of FA binding site are critical for determining the substrate specificity of OleT_MC_

To identify the critical residues of the binding pocket that might affect the substrate specificity of OleT_MC_, we first performed the alanine scan using the “alanine scan” tool in MOE. We scanned a set of 21 residues located at the FA docking sites of FA-bound OleT_MC_ complexes (Supplementary Figure S5A) by substituting each candidate with Ala residue. Regardless of C10:0-C18:0 FAs, we found that the mutation at P72A or I171A increased the stability, while F174A or F296A decreased the stability (Figure 3D). In contrast, the mutation at I177A or V294A showed different stabilities for different FAs. Based on this first round of alanine scan and stability analysis, we narrowed the initial large set of 21 candidate residues to the 6 promising candidates, including P72, I171, F174, I177, F296 or V294, that might have a significant role for the substrate specificity of OleT_MC_.

Next, we performed a comprehensive residue scan for these 6 candidate residues using the “residue scan” tool in MOE. Specifically, we generated a set of 54 OleT_MC_ variants by systematically substituting each candidate with nine different hydrophobic residues including glycine (Gly, G), Ala, Val, Leu, Ile, Pro, Phe, methionine (Met, M), and tryptophan (Trp, W) and performed the stability analysis (Supplementary Figures S7A-F). By determining the largest Δstability differences between C18:0 FA-bound and C10:0 FA-bound OleT_MC_complexes for each OleT_MC_ variant, we could select a list of the top five mutants, including I177W, P72M, F296W, I177F, and P72F (Supplementary Figure S7G), for detail structure analysis. Interestingly, all these five OleT_MC_ variants, selected by combination of alanine and residue scans, had the mutated residues located at the tail of the FA binding pocket (Figures 4C-4H).

To determine whether the top 5 OleT_MC_ variants are responsible for substrate preferences of OleT_MC_, we next generated their 3D structures in MOE, followed by docking simulation of these variants with various FAs (C10:0-C18:0). Our result shows that the two variants, OleT_MC_ P72M and F296W, significantly shifted substrate preferences while the other variants, OleT_MC_ I177W, I177F, and P72F, did not (Figure 4B). For the OleT_MC_ F296W model, we observed that the correct docking poses, whose Arg246 should interact with the carboxylic functional group of FAs via hydrogen bonding, were no longer detected with C16:0 and C18:0 FAs. It has the lowest ΔG_bind_ of −8.43 kcal/mol with C14:0 FA and the highest ΔG_bind_ of −6.72 kcal/mol with C10:0 FA. Likewise, OleT_MC_ P72M showed the lowest ΔG_bind_ of −8.23 kcal/mol with C14:0 FA and the highest ΔG_bind_ of −6.18 kcal/mol with C18:0 FA. These results suggest that P72M and F296W variants can shift the substrate preferences from longer to shorter FAs.

#### 2.3.4 Reconfiguration of the binding pocket is responsible for shifting the substrate preference in OleT_MC_ variants

To elucidate the underlying mechanism for shifting the substrate preference of OleT_MC_, we compared the structures of OleT_MC_ variants, including I177W (Figure 4D), P72M (Figure 4E), F296W (Figure 4F), I177F (Figure 4G), and P72F (Figure 4H), and its wildtype (Figure 4C). Our result shows that the disruption of the FA binding pockets affected substrate preferences of OleT_MC_ P72M (Figure 4E) and F296W (Figure 4F). Specifically, they interfered with the access and docking of the longer C16:0-C18:0 FAs. Furthermore, since OleT_MC_ F296W could not dock C16:0 and C18:0 FAs but OleT_MC_ F72M could, it implies that the size of mutated residue (e.g. Trp (W) having a larger size than Met (M)) can play a significant role in changing the substrate specificity of OleT_MC_.

Taken altogether, the residues at the tail of FA binding pocket of OleT_MC_, such as P72 and F296, are critical for determining the substrate specificity of OleT_MC_. Mutating these residues, instead of those at the catalytic site, can provide a promising protein engineering strategy to shift substrate specificity of OleT_MC_.

### 2.4 Establishing the terminal alkene biosynthesis pathway in *E. coli*

We designed the heterologous terminal alkene biosynthesis pathway in *E. coli* BL21 (λDE3), consisting of two genes − the leaderless *tesA\** gene encoding a thioesterase to convert acyl ACPs to FAs and the OleT_MC_ gene encoding a decarboxylase to convert these FAs to terminal alkenes. We chose TesA* because it has higher specificities towards C16:0-C18:0 acyl ACPs than C12:0-C14:0 acyl ACPs (25) to produce corresponding FAs that are preferable substrates for OleT_MC_. Figure 5A-C shows kinetics of cell growth, sugar consumption, and product formation in shake flask experiments of the recombinant *E. coli* EcNN101 engineered to carry the terminal alkene biosynthesis pathway.

**Figure 5:**
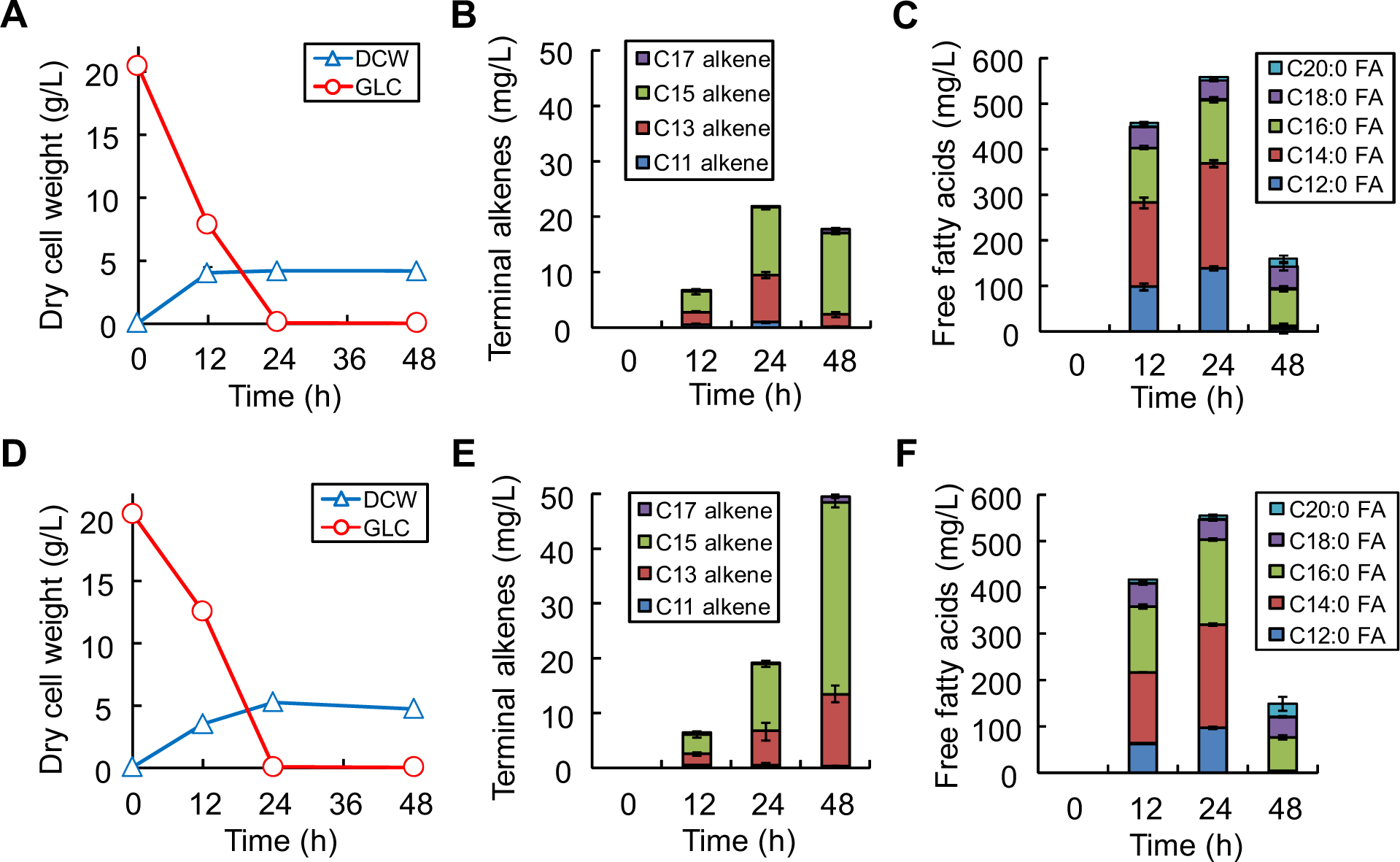
Profiles of alkenes production in *E. coli* (**A-C**) EcNN101 and (**E-F**) and EcNN201. (**A, D**) Cell growth and glucose consumption, (**B, E**) Terminal alkene production, and (**C, F**) FA production.

During the first 24 h of growth phase, EcNN101 could grow and produce odd terminal alkenes. At 24 h, cells completely consumed 20 g/L of glucose and entered the stationary phase with a biomass titer of 4.16 ± 0.07 g/L (Figure 5A). Terminal alkene production peaked at a titer of 21.92 ± 0.69 mg/L, comprised of 4.5% ± 0.2% C11 alkene, 38.9% ± 1.1% C13 alkene, 55.4% ± 0.8% C15 alkene, and 1.2% ± 0.5% C17 alkene (Figure 5B). Besides alkenes, the corresponding FAs were also produced at a much higher titer of 558.03 ± 18.95 mg/L, consisting of 24.9% ± 0.1% C12:0 FA, 41.0% ± 0.2% C14:0 FA, 25.2% ± 0.4% C16:0 FA, 7.6% ± 0.3% C18:0 FA, and 1.4% ± 0.1% C20:0 FA (Figure 5C). The composition of these FAs correlated well with the specificity of TesA* (25). The relatively high FA production clearly implied that OleT_MC_ was the rate-limiting step of the engineered terminal alkene biosynthesis pathway. The result also shows that the C15 alkene was produced at the highest level even though the fraction of C16:0 FA was lower than that of C14:0 FA and relatively similar to that of C12:0 FA. This result is consistent with the substrate preference of OleT_MC_ towards C16:0 FA characterized *in vitro* and also implies that C16:0 FA was not limiting for decarboxylation. The production of C17 terminal alkene, however, was relatively low likely due to the low availability of C18:0 FA.

During the stationary phase (after 24 h), no glucose was available and about 398.15 ±4.79 mg/L saturated FAs were consumed primarily for cell maintenance while cell concentration remained relatively constant. At 48 h, the alkene titer was slightly decreased to 17.78 ± 0.63 mg/L probably due to cell lysis and/or evaporation. The final alkene yield was 0.87 ± 0.03 mg/g. It is interesting to notice that EcNN101 did not produce terminal alkenes during stationary phase even though degradation of saturated FAs, highly reduced substrates, could generate available NAD(P)H for FA decarboxylation via the -oxidation pathway. This result implies that olefin production might be limited by the efficiency of electrons transferred to OleT_MC_ for decarboxylation.

### 2.5 Improving terminal alkene production by enhancing electron shuttling to OleT_MC_

It is known that the electron flow from NAD(P)H to the terminal P450 enzyme is facilitated by a two-component redox system such as ferredoxin reductase (FDR) and NAD(P)H-dependent ferredoxin (FDX). Most of the bacterial P450s belonging to the class I P450s use this two-component redox system for shuttling electrons (26, 27). We hypothesized that the OleT_MC_activity in EcNN101 might have been limiting during the stationary phase due to the lack of a two-component redox system. To test this hypothesis, we constructed EcNN201 that contains both the terminal alkene biosynthesis pathway and a two-component redox system well-characterized for *E. coli*. This redox system consists of an NAD(P)H-dependent putidaredoxin reductase (CamA) and a [2Fe–2S] putidaredoxin (CamB) transferring two electrons, one at a time, from NAD(P)H to the P450 enzyme (26–28).

Like EcNN101, EcNN201 could produce terminal alkenes during the growth phase (Figure 5E). At 24 h, EcNN201 reached a cell concentration of 5.29 ± 0.05 g/L and entered the stationary phase after completely consuming 20 g/L glucose (Figure 5D). EcNN201 produced 19.25 ± 2.03 mg/L terminal alkenes, comparable to EcNN101. The composition of terminal alkenes produced by EcNN201 comprised of 2.4% ± 1.5% C11 alkene, 32.1% ± 5.8% C13 alkene, 64.3% ± 6.7% C15 alkene, and 1.2% ± 0.3% C17 alkene. Like EcNN101, EcNN201 produced a high amount of saturated FAs (556.12 ± 4.44 mg/L) consisting of 17.3% ± 0.3% C12:0 FA, 40.0% ± 0.5% C14:0 FA, 33.3% ± 0.3% C16:0 FA, 7.8% ± 0.4% C18:0 FA, and 1.6% ± 0.3% C20:0 FA (Figure 5F). Overall, the terminal alkene production phenotypes were similar between EcNN101 and EcNN201 during the growth phase. This result implies that reducing equivalents were primarily channeled for ATP generation and biomass synthesis that are thermodynamically favorable under aerobic conditions, and hence likely became limited for decarboxylation to produce target alkenes.

However, during the stationary phase where glucose was not available, EcNN201 consumed a total amount of 407.45 ± 22.19 mg/L FAs for not only cell maintenance but also terminal alkene production. The terminal alkene production was increased up to 58% higher during the stationary phase than the growth phase, underlying the critical functional role of the redox system responsible for enhanced terminal alkene production. EcNN201 produced up to 49.64 ± 1.33 mg/L terminal alkenes, consisting of 0.7% ± 0.1% C11 alkene, 26.4% ± 2.6% C13 alkene, 70.7% ± 3.0% C15 alkene, and 2.2% ± 0.4% C17 alkene. At 48 h, the alkene titer and yield were 49.64 ± 1.33 mg/L and 2.44 ± 0.06 mg/g, respectively. In comparison to EcNN101, the terminal alkene production of EcNN201 increased by 2.8 fold. It is interesting to observe that C12:0-C14:0 FAs were primarily degraded for cell maintenance and C16:0 FA utilized for alkene biosynthesis based on the FA distributions at 24 and 48 h (Figure 5F). This degradation phenotype is consistent with the distribution of terminal alkenes as well as the substrate preference of the endogenous FA synthetase FadD of *E. coli* responsible for catalyzing the first step of the β-oxidation (29).

The FA decarboxylation was clearly the rate-limiting step in our engineered terminal alkene biosynthesis pathway due to high accumulation of saturated FAs observed during the growth phase. The main cause of limited decarboxylation is inefficient electron transfer. We can observe that EcNN101 could not produce terminal alkenes during this stationary phase due to lack of the electron transfer to OleT_MC_. This limitation could be overcome by introducing the two-component redox system in EcNN201. Specifically, the FA degradation during the stationary phase in EcNN201 generated high levels of reducing equivalents (i.e. NAD(P)H) via β-oxidation, and the redox system helped channel electrons to OleT_MC_, thereby improving terminal alkene production.

EcNN201 produced terminal alkenes at a very comparable level to the recombinant *E. coli* harnessing the OleT_JE_ decarboxylase from *Jeotgalicoccus sp.* (16). In our study, we did not observe EcNN101 and EcNN201 producing alkadienes even though unsaturated FAs were produced (Supplementary Figure S8). Because the current terminal alkene production in EcNN201 is very inefficient, improving carbon and electron fluxes via metabolic engineering is critical for enhanced terminal alkene production in future study. In addition, controlling environmental conditions (e.g., sufficient supply of oxygen and use of highly reduced substrates) can potentially help improve terminal alkene production.

## 3. CONCLUSION

In this study, we discovered and characterized a novel P450 FA decarboxylase OleT_MC_ for H_2_O_2_-independent biosynthesis of odd-chain, terminal alkenes. By combining the homology modeling, *in silico* residue mutation analysis, and docking simulation with direct experimental evidence, we elucidated the underlying mechanism that determines the substrate preferences of OleT_MC_. In addition, we demonstrated the direct biosynthesis of medium‐ and long-chain terminal alkenes in an engineered *E. coli* from fermentable sugars, abundant from lignocellulosic biomass. We found that the inefficient electron transfer in OleT_MC_ was the rate limiting step that could be overcome by introducing a two-component redox system. Our results help lay out a foundation for future study to modulate fatty acid thioesterases and Ole_TM_ specificities to produce designer terminal alkenes with desirable carbon chain characteristics. Overall, this study provides a better understanding of the novel functions of FA decarboxylases and helps guide future protein and metabolic engineering for enhanced terminal alkene production in a recombinant host.

## 4. MATERIALS AND METHODS

### 4.1 Bacterial strains and plasmids

Table 1 shows a list of bacterial strains and plasmids used in this study. *E. coli* TOP10 was used for molecular cloning while BL21 (λDE3) was employed as an expression and characterization host. All plasmids were constructed by using a modified pETite* (30), a derivative of pETite C-His backbone vector (Lucigen Corp., WI, USA), suitable for the BglBricks gene assembly method (31). Primers used to construct the plasmids in this study are listed in Table 2.

**Table 1:** List of strains and plasmids.

| Strains/plasmids | Genotype | Sources |
| --- | --- | --- |
| *Strains* |  |  |
| *P. putida* | wildtype | ATCC17453 |
| *M. caseolyticus* | wildtype | ATCC13548 |
| TOP10 | F<sup>-</sup> mcrA Δ(mrr-hsdRMS-mcrBC) φ80lacZΔM15<br>ΔlacX74 nupG recA1 araD139 Δ(ara-leu)7697<br>galE15 galK16 rpsL(Str<sup>R</sup>) endA1 λ<sup>-</sup> | Invitrogen |
| BL21 (λDE3) | F<sup>-</sup> ompT hsdS<sub>B</sub> (r<sub>B</sub><sup>-</sup> m<sub>B</sub><sup>-</sup>) gal dcm (λDE3) | Invitrogen |
| EcNN101 | BL21 (λDE3) carrying pNN32 | this study |
| EcNN201 | BL21 (λDE3) carrying pNN32 and pNN34 | this study |
| *Plasmids* |  |  |
| pETite C-His | pBR322 *ori*; kan<sup>R</sup> | Lucigen |
| pETite* | kan<sup>R</sup> | Layton 2014 |
| pCT71 | pETite* p<sub>T7</sub>:: *tesA* *::T<sub>T7</sub>; kan<sup>R</sup> | this study |
| pNN33 | pETite* p<sub>T7</sub>::*oleT<sub>MC</sub>*::T<sub>T7</sub>; kan<sup>R</sup> | this study |
| pNN32 | pETite* p<sub>T7</sub>:: *tesA* *:: *oleT<sub>MC</sub>*:: T<sub>T7</sub>; kan<sup>R</sup> | this study |
| pNN34 | pETite* p<sub>T7</sub>::*camAB*::T<sub>T7</sub>; amp<sup>R</sup> | this study |

**Table 2:**
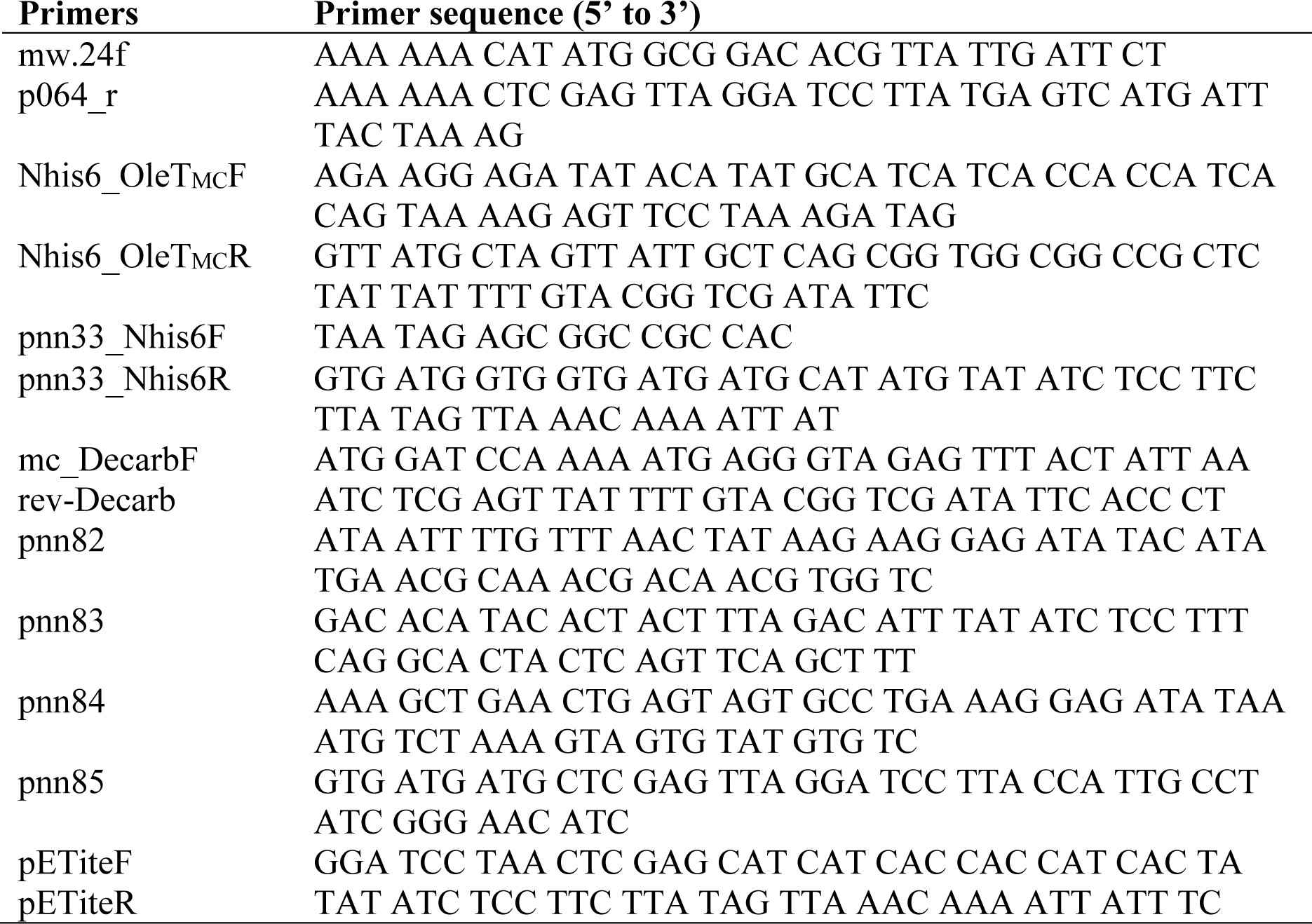
List of primers.

To construct the plasmid pCT71, the leaderless *tesA\** was amplified from the genomic DNA of *E. coli* MG1655 using the primers mw.24f/p064_r, and inserted into pETite* via the NdeI/XhoI sites. The gene *tesA*\* was derived from *tesA* whose leader peptide sequence was removed to keep the encoded protein TesA* localized in the cytosol (32). The plasmid pNN33 was constructed by the Gibson gene assembly method (33) using 2 DNA fragments: i) the decarboxylase gene OleT_MC_ amplified from the genomic DNA of *M. caseolyticus* using the primers Nhis6_OleT_MC_F/Nhis6_OleT_MC_R and ii) the backbone fragment amplified from pETite* using the primers pnn33_Nhis6F/pnn33_Nhis6R. To construct the plasmid pNN32, OleT_MC_ was amplified using the primers mc_decarbF/rev_decarb and inserted into pCT71 via the BamHI/XhoI sites. The plasmid pNN34 was constructed by the Gibson gene assembly method using 3 DNA fragments: i) *camA* gene amplified from the genomic DNA of *P. putida* using the primers pnn82/pnn83, ii) *camB* amplified from the genomic DNA of *P. putida* using the primers pnn84/pnn85, and iii) the backbone fragment amplified from pETite* using the primers pETiteF/pETiteR. All plasmid constructs were confirmed by enzyme digestion, PCR amplification of the respective genes, and sequencing.

The strain EcNN101 was generated by introducing pNN32 into BL21 (λDE3) via electroporation. Similarly, the strain EcNN201 was created by co-transforming pNN32 and pNN34 into BL21 (λDE3).

### 4.2 Medium and strain characterization

#### Medium

For molecular cloning and protein expression, Luria-Bertani (LB) medium containing 5 g/L yeast extract, 10 g/L tryptone, 5 g/L NaCl, and antibiotics (if applicable) was used for *E. coli* and *P. putida* cultures. The medium for cultivating *M. caseolyticus* was comprised of 5 g/L glucose, 5 g/L yeast extract, 10 g/L casein peptone, and 5 g/L NaCl. The hybrid M9 medium used for strain characterization contained 1X M9 salt solution (34), 1 mM MgSO_4_, 10 mM CaCl_2_, 1 mL/L stock trace metals solution, 5 g/L yeast extract, 20 g/L glucose, and appropriate antibiotics (e.g., 50 μg/mL kanamycin or 100 μg/mL ampicillin for single plasmid selection or 25 μg/mL kanamycin plus 50 μg/mL ampicillin for double plasmid selection) (35).

#### Strain characterization

For terminal alkene production experiments, single colonies were inoculated in 15 mL tubes containing 8 mL of LB medium and incubated in a rotary shaker at 37°C with a shaking rate of 175 rpm for 12 hour (h). The overnight seed culture was then transferred into the hybrid M9 medium with an initial OD_600nm_ ∼0.05 in 500 mL baffled shake flasks with a 100 mL working volume. When reaching the exponential phase of OD_600nm_∼0.6, the cell culture was first supplemented with 1 mM 5-aminolevulinic acid (ALA) to enhance the yield of active P450 (36), followed by IPTG induction at a working concentration of 0.5 mM, and run for a total of 48 h. Samples were collected for determining cell growth, substrate consumption, and product formation. All experiments were performed in biological triplicates.

### 4.3 Enzyme expression, purification, and characterization

BL21 (λDE3) pNN33 was used to express OleT_MC_ His-tagged at the N-terminus in LB medium at 20 celcius (°C). Exponentially grown cells (OD_600nm_ ∼0.6) were induced with IPTG at a working concentration of 0.5 mM. After 20 h, cells were collected, washed, and resuspended in 50 mM phosphate buffer (pH 7.5). The resuspended cells were disrupted by ultrasonication with a sonic dismembrator (Model # FB120, Thermo Fisher Scientific Inc., MA, USA) and then centrifuged at 13,500 rcf for 20 min at 4°C to collect the soluble crude cell extract for downstream protein purification. The sonication protocol was set with 70% amplitude with cycles of 5 second ON/10 second OFF pulses on ice for 15 minutes.

The expressed OleT_MC_ protein was semi-purified by the HisPur Ni-NTA Spin column (cat # PI88224, Thermo Fisher Scientific, MA, USA) according to the manufacturer's instruction. Following incubation for 1 h at 4°C, the resin was washed three times with wash buffer (20 mM of imidazole) and the His-tagged OleT_MC_ was eluted from the resin by adding elution buffer (500 mM of imidazole). Then, the final protein sample was concentrated using Amicon Ultra centrifugal filters (cat # UFC801024, Merck, NJ, USA). The purified protein was analyzed by the 12% SDS-PAGE and the protein concentration was determined by the Bradford assay (37).

To quickly test the active form of OleT_MC_ (i.e., the actual heme content of P450 protein), the spectrophotometric carbon monoxide (CO) difference spectral analysis was routinely carried out to determine the maximum characteristic absorbance at 450 nm (38). First, 0.5 mL of 20 mg/mL purified protein sample was diluted in 4.5 mL of 50 mM Tris-HCl buffer (pH 7.6) containing 1 mM EDTA and 10% (v/v) glycerol. The solution was then supplemented with a few crystals of sodium dithionite and saturated with ∼30-40 bubbles of CO at a rate of 1 bubble per second. The maximum spectrophotometric absorbance peak at 450 nm indicates the reduced P450 complexed with CO and hence confirms the enzymatic catalytic center of OleT_MC_ is active.

The H_2_O_2_-independent FA decarboxylase activity of OleT_MC_ was characterized *in vitro* with various linear, saturated C8:0-C20:0 FAs. Each reaction assay has a working volume of 500 μL that contained 1 mM NADH, 10 μM spinach ferredoxin (cat. # F3013, Sigma Aldrich, CA, USA), 0.5 unit spinach ferredoxin reductase (cat. # F0628, Sigma Aldrich, CA, USA), 20μg/mL purified OleT_MC_, and 0.05 mM FAs. The assay was conducted at 37°C for 60 min. After the reaction, alkenes were extracted with ethyl acetate, and analyzed by gas chromatography coupled with mass spectrometry.

### 4.4 Analytical methods

#### Cell growth

Cell optical density was measured at OD_600nm_ using a spectrophotometer (Spectronic 20+, Thermo Fisher Scientific, MA, USA). The correlation between the cell optical density and dry cell weight (DCW) was determined to be 1 OD_600nm =_0.48 g DCW/L.

#### High performance liquid chromatography (HPLC)

The HPLC Shimadzu system equipped with a BioRad Aminex HPX 87-H column (cat # 1250140, BioRad, CA, USA) and both RID and UV-VIS detectors (Shimadzu Scientific Instruments, Inc., MD, USA) were used to quantify extracellular metabolites (e.g., glucose, organic acids, and alcohols). The running method used 10 mN H_2_SO_4_ as a mobile phase operated at a flow rate of 0.6 mL/min and an oven temperature set at 48°C (39).

#### Gas chromatography coupled with mass spectroscopy (GC/MS)

For FAs analysis, sample preparation and GC/MS methods were described previously (40). For terminal alkene analysis, 500 μL of samples (cells plus supernatants) were transferred to a 2 ml polypropylene microcentrifuge tube with a screw cap containing 100-200 mg of glass beads (0.25-0.30 mm in diameter), 60 μL of 6 N HCl, and 500 μL of ethyl acetate solution containing 1 mg/L of ethyl pentadecanoate as an internal standard. The cells were lysed by bead bashing for 4 min using a Biospec Mini BeadBeater 16 and then centrifuged at 13,300g for 2 min. The extractants of the organic layer were used for the GC/MS analysis. All alkenes were analyzed by using HP6890 GC/MS system equipped with a 30 m x 0.25 mm i.d. 0.25 μm film thickness column plus an attached 10 m guard column (Zebron ZB-5, Phenomenex Inc.) and a HP 5973 mass selective detector. An electron ionization in scan mode (50 to 650 m/z) method was deployed to analyze 1 μL of samples. The column temperature was initially held at 50°C for 1 min, increased by 20°C/min until 300°C, and held for 2 min. Helium was used as a carrier gas and run at 1 mL/min. The mass transfer line and ion source were set at 250°C and 200°C, respectively.

### 4.5 Bioinformatics

For sequence alignment and phylogenetic analysis, protein sequences were retrieved from the National Center for Biotechnology Information (NCBI), and were inputted into MEGA7 (41) and aligned via MUSCLE (42). The phylogenetic tree was generated using the neighbor-joining algorithm with a 1000 bootstrap value. BLASTp was used to calculate the identity of sequences (43) using OleT_JE_ as the template.

### 4.6 Homology modeling and *in silico* analysis of OleT_MC_

#### Homology modeling

The homology model of OleT_MC_ was generated using the version 2015.10 Molecular Operating Environment (MOE) software (21). To obtain the heme-bound protein structure, the reference substrate, heme, was first extracted from the best hit, substrate-bound crystallographic structure of OleT_JE_ (PDB:4L40) and then added to the homology model of OleT_MC_. Next, the heme-bound model of OleT_MC_ generated was optimized by energy minimization using the Amber10:EHT (Extended Huckel Theory) force field (44, 45). The Ramachandran plot analysis was performed to determine the overall stereochemical property of the protein model.

#### Docking simulation

To dock various FAs with the heme-bound homology model of OleT_MC_, the 3D structures of various C10:0-C18:0 FAs were first generated by modifying C20:0 FA extracted from OleT_JE_ (PDB:4L40) with the ‘3D builder’ tool of MOE and then optimized by energy minimization using the Amber10:EHT force field. Next, the “site finder” tool of MOE was used to search for potential binding sites. Upon identifying the site consistent with the reported catalytic site of OleT_JE_ (20), dummy atoms were generated to mark potential binding sites. To select the exclusive potential binding site of FA, we also removed some dummy atoms located near heme. Finally, we added the target C10:0-C18:0 FAs to the binding site of the heme-bound homology model of OleT_MC_. All structures were protonated using the “protonate3D” tool of MOE prior to docking simulation.

Docking simulations were carried out as previously described (46). In brief, the induced fit docking protocol employed the Triangle Matcher placement method and the London ΔG scoring function. In our docking simulations, we performed 30 docking interactions for each FA substrate. The binding free energy (ΔG*_bind_*, in kcal/mol) for each binding pose was then minimized using the Amber10:EHT force field and rescored with the GBVI/WSA ΔG scoring function (44). The best scored pose, exhibiting the crucial interaction between the residue Arg246 and carboxylic functional group of the substrate via hydrogen bonding (20) at root-mean-square-deviation (RMSD) < 2 Å, was selected for further analysis. The “surface and maps” tool of MOE was employed to visualize the molecular surface of atoms in the potential FA binding site.

#### In silico mutation analysis

The “alanine scan” and the “residue scan” tools in MOE were used for *in silico* mutation analysis of FA-OleT_MC_ complexes. Specifically, the alanine scanning technique (47, 48) was employed to determine the importance of a specific residue to the stability, affinity, and/or property of the FA-OleT_MC_ complexes upon being substituted with Ala in the binding pocket. Residue scanning technique, also known as site-directed mutagenesis (49), was applied to generate large number of OleT_MC_ variants for the comprehensive mutation study using the selected residues from the alanine scan. By utilizing these tools, we could replace each of the interface residues with a specific residue of interest and calculates the effect of the mutation on the binding free energy (ΔG_bind_) of the complexes. The ΔStability values (kcal/mol) were calculated as the relative binding free energy difference (ΔΔG_bind_) between the mutant (ΔG_mutant_) and wild type (ΔG_wildtype_) in MOE.

## ACKNOWLEDGEMENTS

We would like to thank Dr. Fu-min Menn (Center of Environmental Biotechnology, UTK) for his assistance in developing the GC/MS method, Dr. Jerome Baudry (UTK) for accessing the MOE software, Dr. Donovan Layton (UTK) for assistance with GC/MS and bioinformatics analyses, and members of Trinh lab for proofreading and providing critical comments on the manuscript.

## FUNDING

This research was supported by the laboratory start-up, SEERC, and JDRD seed funds from the University of Tennessee, Knoxville (UTK), a NSF CAREER award (NSF#1553250 to CTT), and a DOE subcontract grant (DE-AC05-00OR22725) by the Center of Bioenergy Innovation (CBI), the U.S. Department of Energy Bioenergy Research Center funded by the Office of Biological and Environmental Research in the DOE Office of Science.

## Conflict of interest statement

None declared

